# PlrA (MSMEG_5223) is an essential polar growth regulator in *Mycobacterium smegmatis*

**DOI:** 10.1101/2022.09.08.507195

**Authors:** Samantha Y. Quintanilla, Neda Habibi Arejan, Parthvi B. Patel, Cara C. Boutte

**Affiliations:** Department of Biology, University of Texas Arlington, Arlington, TX

## Abstract

Mycobacteria expand their cell walls at the cell poles in a manner that is not well described at the molecular level. In this study, we identify a new factor, PlrA, involved in restricting peptidoglycan metabolism to the cell poles in *Mycobacterium smegmatis*. We show that PlrA localizes to the pole tips, and we identify its essential domain. We show that depletion of *plrA* pheno-copies depletion of polar growth factor Wag31, and that PlrA is involved in regulating polar peptidoglycan metabolism and the structure of the Wag31 polar foci.

## Introduction

Expansion of the cell wall is critical for bacterial growth. In rod shaped bacteria, cells expand by elongating the rod, and then divide centrally to propagate daughter cells. Elongation occurs along the lateral walls in many proteobacterial and firmicute species [1]. Polar elongation occurs in several alphaproteobacterial species [2] and in Actinomyecetes [3,4]. In the alphaproteobacterium *Agrobacterium tumefaciens*, polar growth is dependent on Growth Pole Ring (GPR) protein, which forms a ring around the pole and is required for restricting peptidoglycan synthesis to the pole [5,6]. Many Actinobacteria, including mycobacteria, also elongate at the poles [3,7]. Actinobacterial polar growth is dependent on DivIVA-like proteins [8–12], which, like the GPR, restrict peptidoglycan synthesis to the poles [12]. The molecular mechanisms by which GPR and DivIVA proteins mediate polar elongation have not been described.

In Mycobacteria, the polar DivIVA-like protein is called Wag31. Wag31 localizes to the cell poles, with more Wag31 associated with the faster-growing old pole [9,13,14]. While it is clear that Wag31 is essential for establishing the pole and restricting peptidoglycan metabolism to the pole [12,15], it is not at all clear how it works. Wag31 has no enzymatic domains and is cytoplasmic. In firmicutes, DivIVA proteins have been shown to recruit and activate other proteins involved in cell wall synthesis and regulation [16–20]. It is presumed that Wag31 somehow regulates polar peptidoglycan synthesis enzymes. However, despite being immunoprecipitated to find interaction partners in several studies [21–23], Wag31 has never been shown to interact with any other polar peptidoglycan synthesis enzymes.

The complex of cytoplasmic, transmembrane and periplasmic regulators and cell wall enzymes that collectively mediate the ordered elongation of the cell wall is called the elongasome. In lateral growers, the elongasome comprises cytoplasmic regulators like MreB, and peptidoglycan enzymes including RodA and PBP2 [1,24]. These proteins all function together as a complex to allow the ordered insertion of new peptidoglycan. Wag31 has been called an elongasome protein in Mycobacteria [4]; however, it is not at all clear that Mycobacterial elongation is mediated by a large protein complex that functions similarly to the elongasome characterized in *E. coli* and other lateral growers. First, recent work shows that the critical peptidoglycan synthases required for polar growth are not even localized to the pole, but instead are distributed nearly evenly around the cell membrane [25]. There must therefore be a system to activate these proteins only near the pole. One model is that cell wall synthesis is activated by the availability of cell wall precursors such as lipidII. Cell wall precursor enzymes are localized largely to the Intracellular Membrane Domain, a biochemically distinct region of the inner membrane that is localized mostly peri-polarly [15,26,27]. IMD enzymes, such an MurG are therefore near the pole, but not at the pole, and they do not co-localize with Wag31. Thus, it remains an open question how Wag31 can regulate the activity of enzymes when it does not co-localize either with those enzymes or the production of their substrates.

Because polar growth in mycobacteria is so poorly understood, we reasoned that there are likely many genes involved in this process that have not yet been characterized. In this study, we describe initial characterization of one of those factors. In a previous study, we immunoprecipitated the transmembrane division factor FtsQ from *M. smegmatis* and identified several uncharacterized interactors [28]. One of these was MSMEG_5223 (Rv1111), which we found localized to the cell poles as well as the septum [28]. In this study we show that MSMEG_5223, hereafter called PlrA, is essential for polar elongation in *Msmeg*, and that, like Wag31, it restricts peptidoglycan metabolism to the pole. We also show that only the N-terminus of PlrA is essential. Finally, we show that depletion of PlrA affects the structure of the Wag31 focus at the pole, suggesting that PlrA may regulate Wag31 oligomerization.

## Results

### PlrA is essential for polar peptidoglycan metabolism and elongation

*plrA* (MSMEG_5223, Rv1111) is predicted by TnSeq to be essential for survival in both *Mycobacterium tuberculosis* [29] and *Msmeg* [30]. To study its function genetically, we made a strain, Ptet:: *plrA*, in which it can be transcriptionally depleted by removing the inducer anhydrotetracyline (Atc). We grew the Ptet:: *plrA* strain to logarithmic phase, then washed out the Atc and measured survival using CFUs. Our results (Fig. 1A) show that PlrA is essential for survival. We then examined the *plrA* -depleted cells microscopically and found that after 30 hours of depletion, they are short with bulgy poles (Fig. 1B). These data show that *Msmeg* is unable to elongate properly and is unable to control cell wall structure at the poles without PlrA. We therefore conclude that PlrA is an essential polar elongation factor. Because PlrA does not have a predicted enzymatic domain, we infer that it is a regulator of polar elongation. We therefore name it *plrA* for <u>p</u>o<u>l</u>e <u>r</u>egulator A.

**Figure 1.**
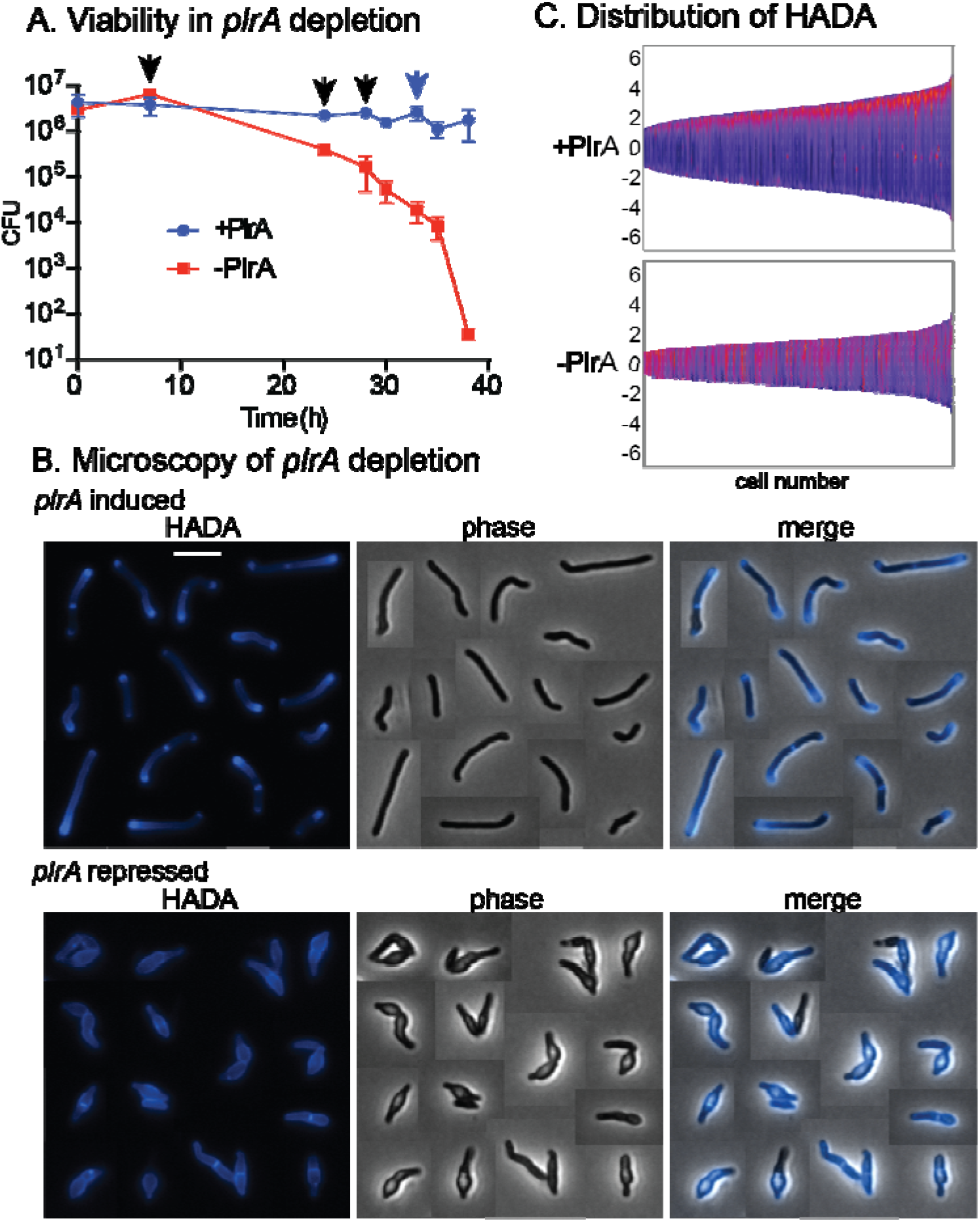
PlrA is an essential polar growth regulator. A) CFU of the Ptet::*plrA* strain in the presence (*plrA* induced) or absence (*plrA* repressed) of inducer Atc. Three independent replicate cultures were used for each condition. Black arrows represent where dilutions were performed in both cultures to prevent them reaching stationary phase. Blue arrow indicates where dilution was performd in the +Atc (+PlrA) culture only. B) Micrographs of a Ptet::*plrA* strain with *plrA* induced (top, +Atc) or repressed (bottom, -Atc), then stained with the fluorescent D-alanine HADA for 15 minutes. Cells from different images were cut and pasted together so that a representative collection of cells could be shown. The scale bar on the top left image is 5 microns, and applies to all images. C) Demographs of HADA intensity along the length of cells (Y axis) in *plrA* induced (top) and depleted (bottom) cells from panel B. Cells were sorted by size (X axis) and pole sorted, so that brightest pole is set at the top. Lighter colors represent higher HADA intensity.

*Mycobacteria* insert new peptidoglycan and other cell wall materials near the cell poles to elongate [7,13]. We used the fluorescent D-alanine HADA [31] to probe how the distribution of peptidoglycan metabolism in the cells was affected by *plrA* depletion. We found that *plrA* depletion led to delocalized HADA staining, instead of the typical poles and septa pattern (Fig. 1BC). HADA reports on both insertion of new peptidoglycan and remodeling of existing peptidoglycan [27,32], so these data cannot tell us whether new peptidoglycan synthesis is occurring all along the lateral walls, or whether peptidoglycan remodeling is just de-regulated. However, because most of the HADA signal comes from peptidoglycan remodeling [32], and because cell elongation is clearly slowed, we conclude that *plrA* likely promotes polar-adjacent peptidoglycan remodeling, as well as polar insertion of new peptidoglycan. Short, bulgy cells and delocalized peptidoglycan metabolism is also seen when the essential DivIVA homolog Wag31 is depleted [9,12].

### PlrA localizes to the tips of both cell poles

In our previous study, we showed that PlrA localizes to cell poles and septa in *Msmeg* [28]. The polar growth regulator Wag31 has a similar localization pattern, and is seen to localize more strongly to the faster-growing old pole [23]. We stained cells expressing PlrA-GFPmut3 with HADA, which stains the old pole more brightly [32], in order to see if PlrA localizes in a similar pattern. We found (Fig. 2) that PlrA-GFPmut3 does have slightly brighter signal at the pole with brighter HADA staining, indicating that it localizes more to the faster growing pole. However, on average PlrA localization is similar between the two poles, compared to the significant difference in HADA staining between the new and old poles (Fig. 2BC). These data show that PlrA has a similar localization pattern as Wag31, in that it localizes to the pole tips [9]; however, the minimal asymmetry in PlrA localization suggests that the amount of PlrA at the pole is likely not responsible for regulating the asymmetry of polar elongation [13,33].

**Figure 2.**
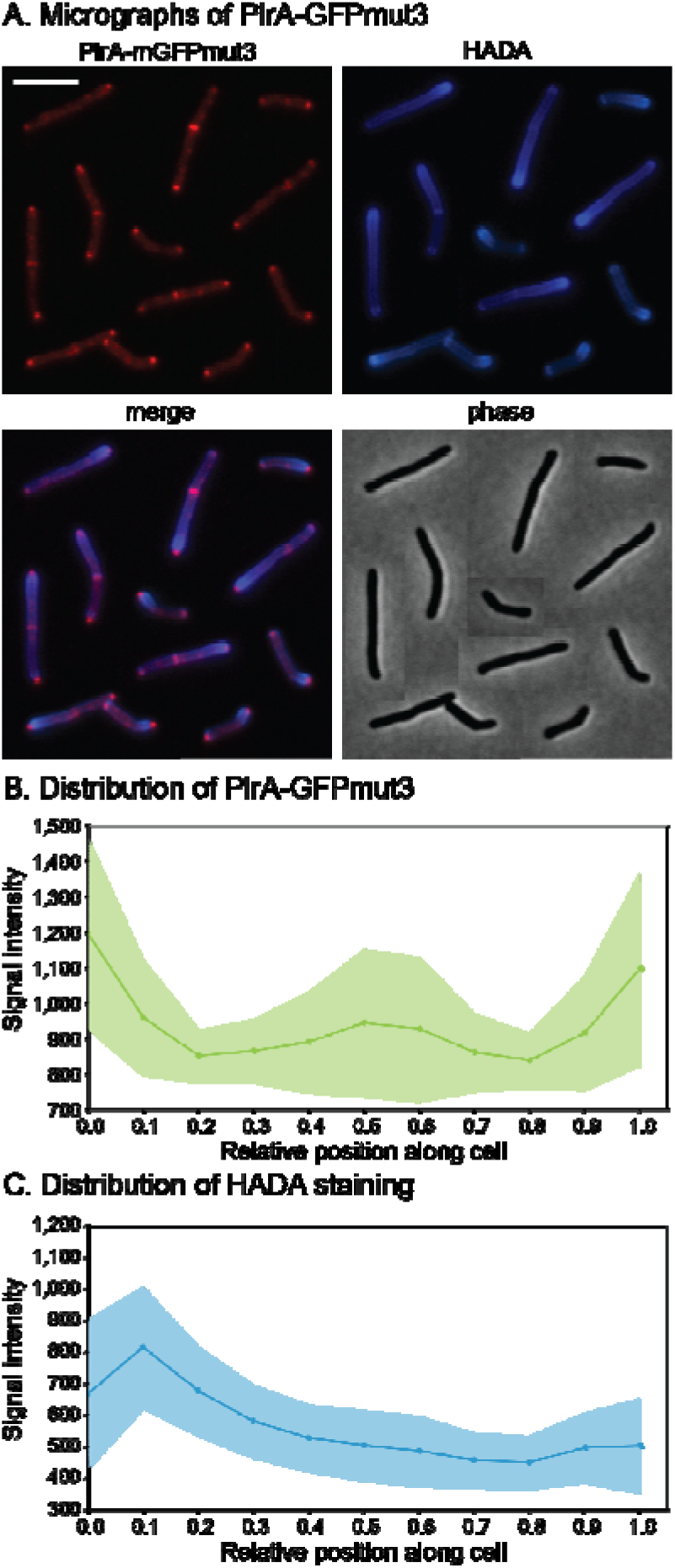
PlrA localizes to the pole tips. A) Micrographs of *Msmeg* mc^2^155 expressing PlrA-GFPmut3 as a merodiploid and stained with the fluorescent D-alanine HADA. The GFP signal was false colored to red to make the channels easier to distinguish. The scale bar on the top left image is 5 microns, and it applies to all images. Cells from different images were cut and pasted together so that a representative collection of cells could be shown. BC) Mean intensity of PlrA-GFPmut3 (B) and HADA (C) signal along the length of ∼300 cells, at least 100 from each of three biological replicates. Center line is the mean signal, lighter area is the standard deviation. Cells were pole sorted, so the brightest pole in the HADA channel is set to 0, and the dimmer pole is set to 1 on the X axis.

### The C-terminus of PlrA is dispensable, while the N-terminus is essential

PlrA has an N-terminal membrane domain with four predicted transmembrane passes and and C-terminal predicted cytoplasmic domain [34]. Because PlrA has no significant sequence similarity to any gene characterized in bacteria, we sought to dissect its essentiality, by determining whether both or only one of these domains was essential. First, we used Consurf [35] to identify the relative conservation of each amino acid in the *Msmeg* protein. This analysis shows that the N-terminal membrane domain is more highly conserved than the C-terminal cytoplasmic domain (Fig. 3A).

**Figure 3.**
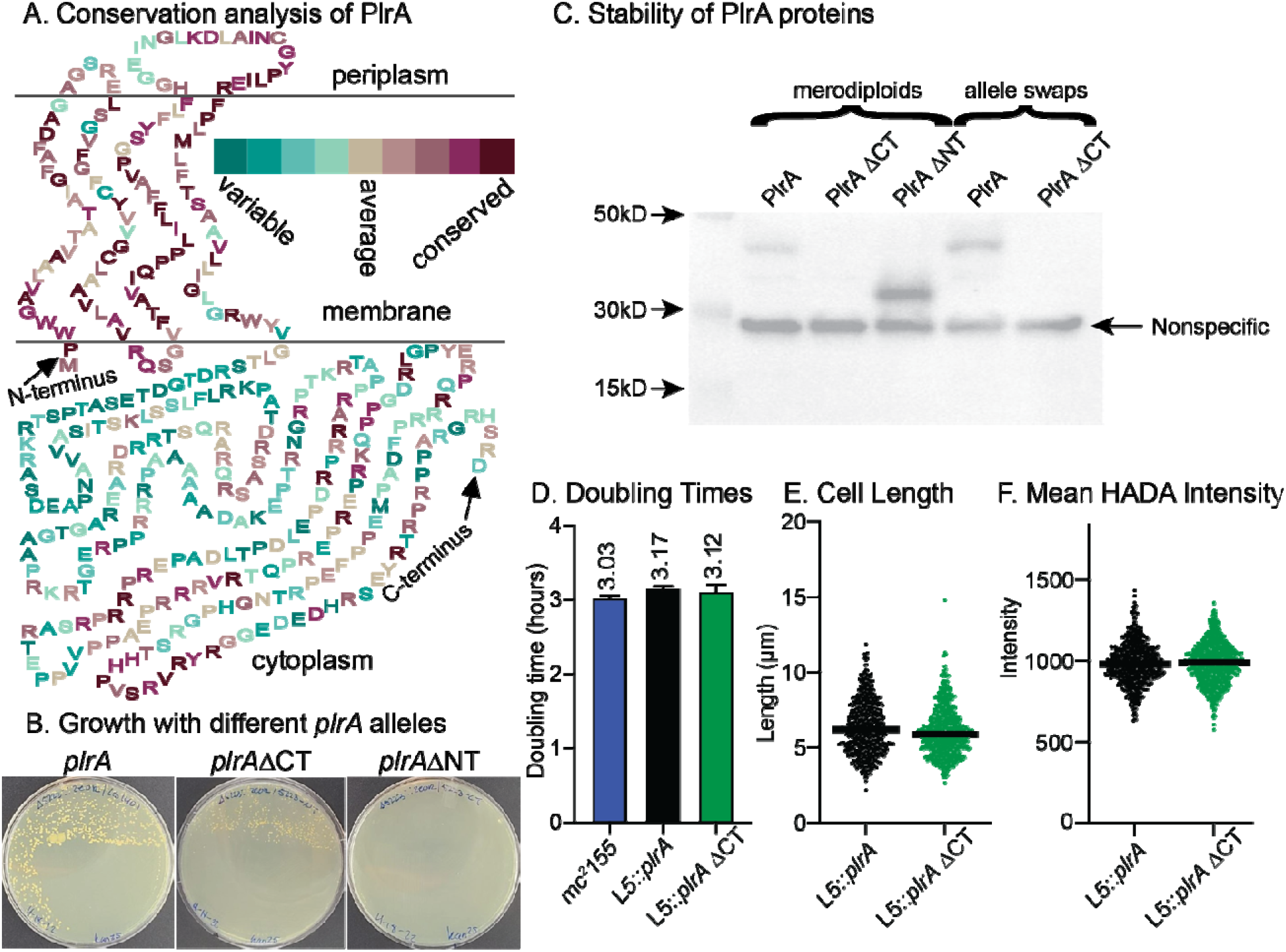
The C-terminal domain of PlrA is dispensable. A) PlrA protein sequence arranged in either periplasmic, membrane or cytoplasm as predicted by TMHMM. Amino acids are colored according to conservation as measured by Consurf analysis. B) Plates resulting from L5 allele swapping of the wild-type *plrA* with full-length *plrA-*strep, or *plrA*ΔCT-strep or *plrA*ΔNT-strep. The experiment was arranged so that the original, wild-type allele is lost in any colonies. C) Western blot of *Msmeg* strains carrying either *plrA-*strep, *plrA*ΔCT-strep and *plrA*ΔNT-strep as merodiploids or allele swaps. PlrA*-*strep is 43kD, PlrAΔCT-strep is 13kD, PlrAΔNT strep is 31kD. D) Doubling times calculated from growth curves in 7H9 of the mc^2^155 parent, Δ*plrA* L5::*plrA-*strep, and Δ*plrA* L5::*plrA*ΔCT*-*strep strains. E) Cell lengths of the Δ*plrA* L5::*plrA-*strep, and Δ*plrA* L5::*plrA*ΔCT*-*strep strains in logarithmic phase, as quantified from phase microscopy images by MicrobeJ analysis. F) Average HADA intensity per cell of cells from E.

Then, we used L5 allele swapping [36] to replace the full-length *plrA* with *plrA*ΔCT (residues 1-117) or with *plrA*ΔNT (residues 118-368). In this method, a copy of *plrA* under the control of a tet-inducible promoter was cloned into a nuoR vector carrying the TetR repressor and inserted into the *Msmeg* genome at the L5 phage integrase site, then the endogenous copy of *plrA* was deleted. We then cloned the full-length and truncation alleles of *plrA*, also under tet-promoters, into a kanR L5 integrating vector without the *tetR* gene. Transformation of these kanR vectors into the *Msmeg* strain carrying *plrA* only at the L5 site could result in either nuoR +kanR double integrants, or in kanR nuoS allele swaps. Because only the original L5 vector carries *tetR*, which will repress expression of either of the *plrA* alleles in either L5 vector, we plate the transformants without the Atc inducer and therefore select against the double integrants. Because *plrA* is essential (Fig. 1A), in this setup, we will only get colonies on the transformation plate if the *plrA* allele in the second, kanR vector is functional enough to support growth. We found that a strain carrying only *plrA*ΔCT is viable, while a strain carrying only *plrA*ΔNT is not viable. This indicates that the more highly conserved N-terminal domain is essential, while the C-terminal domain is not (Fig. 3B). All *plrA* alleles were cloned with a C-terminal strep tag, and we tested the stability of the PlrA truncations by western blot (Fig. 3C). We made merodiploid strains of all the constructs so we could test whether the PlrAΔNT protein is stable. We found that PlrAΔNT is even more stable than the full length protein, while the PlrAΔCT protein is less stable, and did not yield a detectable band on the western blot in either the merodiploid or the allele swap strain. This shows that the C-terminal domain of PlrA is not essential for function, and is not required merely for protein stabilization. These data also suggest that very little PlrA is needed for survival, as the PlrAΔCT protein is undetectable by western, despite supporting growth.

We next tested whether the PlrA C-terminal domain contributes to growth in logarithmic phase. We found that the *plrA*ΔCT strain has no defects in growth rate (Fig. 3D), cell morphology (Fig. 3E) or peptidoglycan metabolism as measured by fluorescent D-amino acid staining (Fig. 3F). These data show that the C-terminal domain of PlrA is entirely dispensable for normal logarithmic phase growth.

### Depletion of PlrA causes atypical accumulation of Wag31 at the poles

Because the *plrA* depletion (Fig. 1) exhibited a similar phenotype as the *wag31* depletion [12], we hypothesized that these two proteins may work together to regulate polar growth. We first sought to determine whether Wag31 localization is dependent on PlrA. We transformed a vector expressing a Wag31-mRFP fusion into the Ptet::*plrA* depletion strain. We grew the resulting strain with or without the Atc inducer, then HADA-stained the cells and examined them microscopically. We found that Wag31-mRFP still localizes to the cell poles in the cells depleted for *plrA* (Fig. 4A). However, we observed that the size and intensity of the Wag31 foci was more variable in the *plrA*-depleted cells (Fig. 4BCD). Many of the *plrA*-depleted cells, especially the shorter cells which are presumably more severely depleted, have unusually bright and large foci, while other cells have very dim Wag31-mRFP foci (Fig. 4B). In the control cells (left side of Fig. 4), HADA and Wag31-RFP intensity are greater at the same cell pole in each cell, which we expect to be the old pole [32]. In the *plrA-*depleted cells, the new pole can be identified in V-snapping cells as the pole at the vertex of the V. We find, in these V-snaps, that the old pole is often dimmer by HADA than the new pole, while the new pole is usually the one bulging. We find that the unusually bright Wag31 foci are often at HADA-dim old poles (Fig. 4AB), and therefore the cell pole that is brighter by HADA is not also brighter by Wag31-RFP (Fig. 4C).

**Figure 4.**
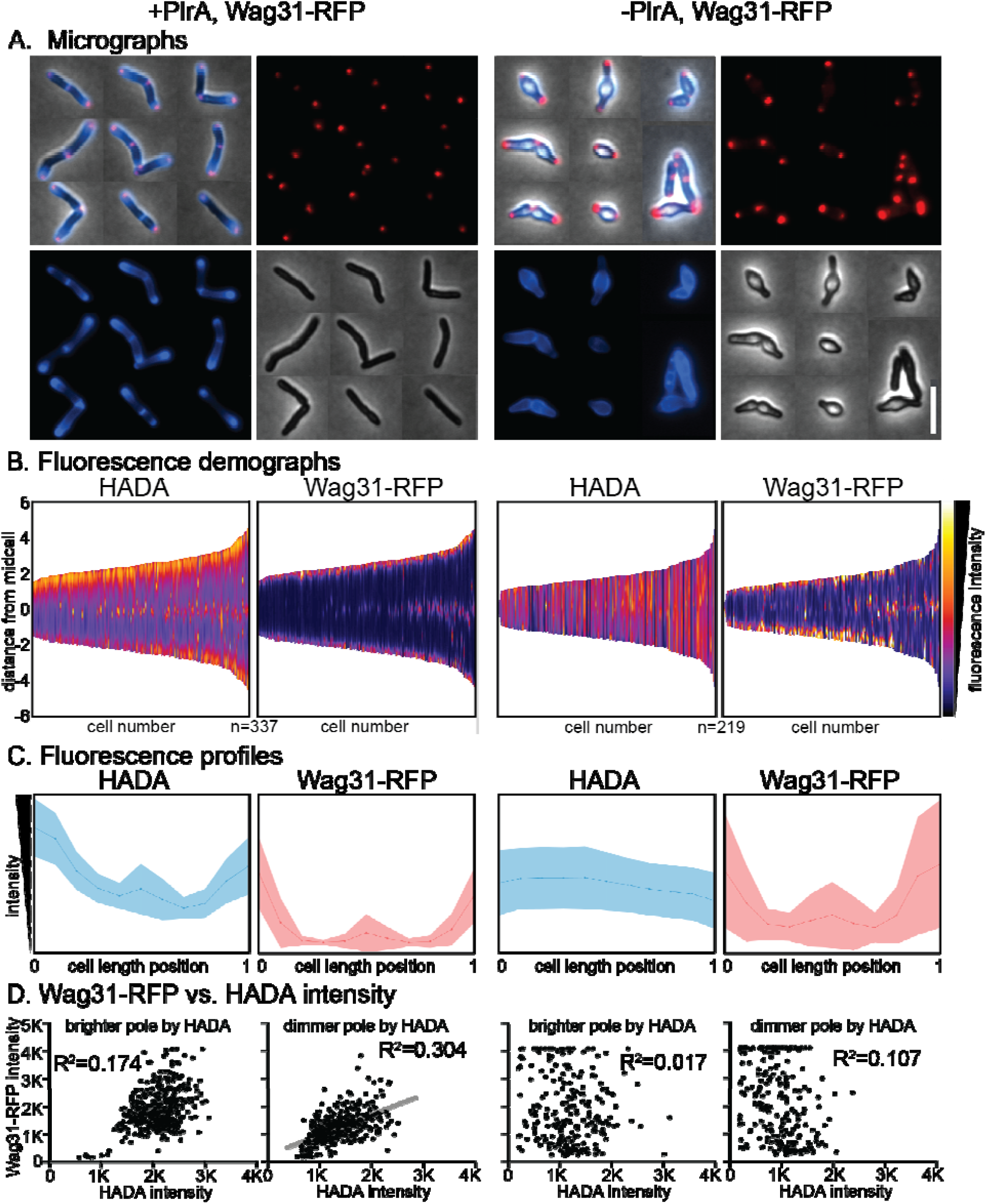
PlrA helps regulate the Wag31 polar foci. A) Micrographs of Ptet::*plrA* Wag31-mRFP strains induced (left) and depleted (right) for *plrA*, and stained with HADA. Blue= HADA fluorescence image. Red= Wag31-RFP fluorescence image. Scale bar on bottom right is 5 microns and applies to all images. B) Demographs of fluorescence intensity of the cell populations imaged in (A). *plrA* induced cells are on the left, *plrA* depleted cells on the right. Cells are arranged shortest to longest along the X axis, and arranged so the pole with the brighter HADA signal is positioned at the top. C) Mean fluorescence intensities (Yaxis) of all the cells from (A,B) at 11 points along the length of each cell. *plrA* induced cells are on the left, *plrA* depleted cells on the right. Darker line in the center is the mean, and shaded area is the standard deviation. Cells are sorted so that the pole with the brighter HADA intensity is set to 0 on the X axis. Both HADA graphs have intensity values between 0-2400 on the Y axis. Both RFP graphs have intensity values between 0-3200 on the Y axis. D) Maximum Wag31-RFP signal (Yaxis) plotted agains the maximum HADA signal (X axis) at each cell pole. *plrA* induced cells are on the left, *plrA* depleted cells on the right. R^2^ values were calculated by linear regression analysis. The gray line is the linear fit on the only graph with a correlation.

To probe the relationship between peptidoglycan metabolism - as measured by HADA staining - and Wag31-RFP localization, we plotted the maximum values of fluorescence intensity at each cell pole against each other (Fig. 4D). We find that in the control cells, the presumed old poles (brighter by HADA) have roughly gaussian distributions of both Wag31 and HADA signal across the population, and there is not a significant correlation between the signal in these two channels. This suggests that in the control cells, all the old poles are similar with respect to peptidoglycan metabolism and Wag31, which is what we expect since all old poles grow at the same rate [13,37]. There is a weak correlation between Wag31-RFP signal and HADA signal in control cells at the presumed new poles (Fig. 4D). However, this makes sense as the new pole undergoes changes throughout the cell cycle: right after division it does not elongate, and so we see less peptidoglycan metabolism (Fig. 4BD), but as the cells mature, the new pole becomes elongation-competent [37], and we see a corresponding increase in Wag31-RFP signal (Fig. 4BD). In the *plrA*-depleted cells, we see a loss of Wag31-RFP signal intensity clustering in both poles, and the correlation between HADA signal and Wag31-RFP signal at the HADA-dim pole is lost. These data suggest that PlrA helps control the structure of the Wag31 focus, as well as the polarity of peptidoglycan metabolism.

## Discussion

It remains an open question how much the “elongasome” model from lateral growing bacteria should serve as inspiration for developing models for polar growth. In this model, cytoplasmic regulators help control periplasmic cell wall enzymes through trans-membrane protein interactions. In both the alphaproteobacterium *A. tumefaciens* and the actinobacterium *C. glutamicum*, this model seems to hold up to some extent, as the key polar peptidoglycan transglycosylase enzymes are localized at the cell poles [38,39], and in *C. glutamicum* is anchored there through a cytoplasmic regulator [39]. We do wish to note that mis-localization of proteins can occur easily during over-expression, for example [40,41]. The model of the elongasome complex does not seem to apply in *Mycobacterium smegmatis*, where the putative polar growth regulator Wag31 does not co-localize with either peptidoglycan precursor enzymes or transglycosylases [23,25–27,42]. Since polar growth appears to work so differently in Mycobacteria, we reasoned that there must be other essential factors involved in this process, and that characterizing those factors may help establish a new model for Mycobacterial polar growth.

Our work shows that the membrane domain of PlrA is the domain essential for polar growth, and the cytoplasmic domain appears to not have any function during logarithmic phase growth (Fig. 3). Non-enyzmatic membrane proteins involved in cell growth and division can either have roles regulating enzymes in the periplasm [43] or the cytoplasm [44], or they can bind and regulate other factors through their membrane-pass regions [45,46]. The fully functional PlrAΔCT protein has only a four-amino acid cytoplasmic loop, while there is a 22-amino acid periplasmic loop. The most highly conserved residues are in the membrane passes and in a region of the periplasmic loop. We therefore think it most likely that PlrA regulates either a periplasmic enzyme, or another membrane protein through membrane contacts.

What could PlrA be doing to Wag31? Previous work has only shown that increased Wag31 at sites in the cell causes increases in polar growth [47]. Our work shows that a large Wag31 focus can be inactive in polar growth when PlrA is missing. The asymmetry of Wag31 foci at the poles, which correlates with the asymmetry of growth, suggests that the conformation or size of the homo-oligomeric Wag31 network could be involved in regulating polar growth (Fig 4BD).

Our work suggests that PlrA is required for a Wag31 focus to permit polar peptidoglycan synthesis (Fig. 4). Perhaps PlrA helps control the chemical structure or shape of the pole, which may, in turn, affect Wag31 oligomer organization. Depletion data suggests that pole structure is dependent on PlrA, not solely on the presence of the Wag31 oligomer, as the Wag31 oligomer remains in place when the poles bulge due to *plrA* depletion (Fig. 4A).

## Materials and Methods

### Bacterial strains and culture conditions

*M. smegmatis* mc^2^155 was cultured in 7H9 (Becton, Dickinson and Co, Sparks, MD) medium with additives as described [28] or plated on LB Lennox agar. E. coli DH5a, TOP10, or XL1-Blue cells were used for cloning. The antibiotic concentrations used for *M. smegmatis* were: 25 mg/ml kanamycin, 50 mg/ml hygromycin, 20 mg/ml nourseothricin, and 20 mg/ml zeocin. The antibiotic concentrations used for *E. coli* strain were: 50 mg/ml kanamycin, 100 mg/ml hygromycin, 50 mg/ml zeocin, and 40 mg/ml nourseothricin. Anhydrotetracyline was used at between 50 and 250 ng/ml for gene induction or repression.

### Strain construction

Knockout of *plrA* was made by first integrating a copy of the gene at the L5 site [48] in the pMC1s vector with a tet-inducible P750 promoter. The endogenous copies of the gene was then knocked out using double stranded recombineering, as described [49]. Vectors were assembled using Gibson cloning [50], some with the SSB enhancement [51].

### Colony forming unit assay

Clones of the Ptet:: *plrA* strain were grown to logarithmic phase in 7H9 with nourseothricin, zeocin, and 500 ng/mL of anhydrotetracyline (Atc). All cultures were washed to remove Atc, and diluted to OD=0.1, Atc was added to half the cultures and allowed to grow. At the 7 hour time point, both cultures were diluted to OD=0.01. At the 24 hour time point, both cultures were diluted to OD=0.2. At the 28 hour time point, both cultures were diluted to OD=0.1. At the 35 hour time point only the +Atc culture was diluted to OD=0.01. Atc was re-added to the +Atc cultures only during the dilutions. CFU were measured on LB plates with nourseothricin, zeocin and Atc.

### Microscopy and image analysis

Microscopy was performed on living cells immobilized on Hdb-agarose pads. A Nikon Ti-2 widefield epifluorescence microscope with a Photometrics Prime 95B camera and a Plan Apo 100x, 1.45 NA objective was used for imaging. The GFPmut3 images were taken with a 470/40nm excitation filter, a 525/50nm emission filter and a 495nm dichroic mirror. The HADA images were taken using a 350/50nm excitation filter, a 460/50nm emission filter and a 400nm dichroic mirror. The mRFP images were taken with a 560/40nm excitation filter, a 630/70nm emission filter and a 585nm dichroic mirror. All images were processed using NIS Elements software and analyzed using FIJI and MicrobeJ [52].

---

**Supplemental Table 1.**
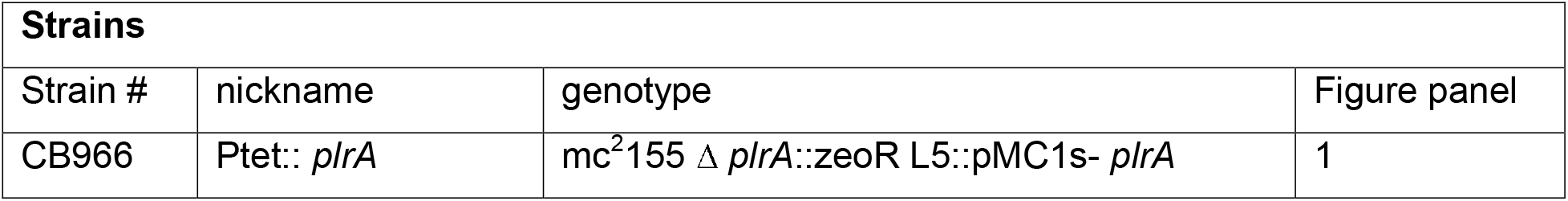
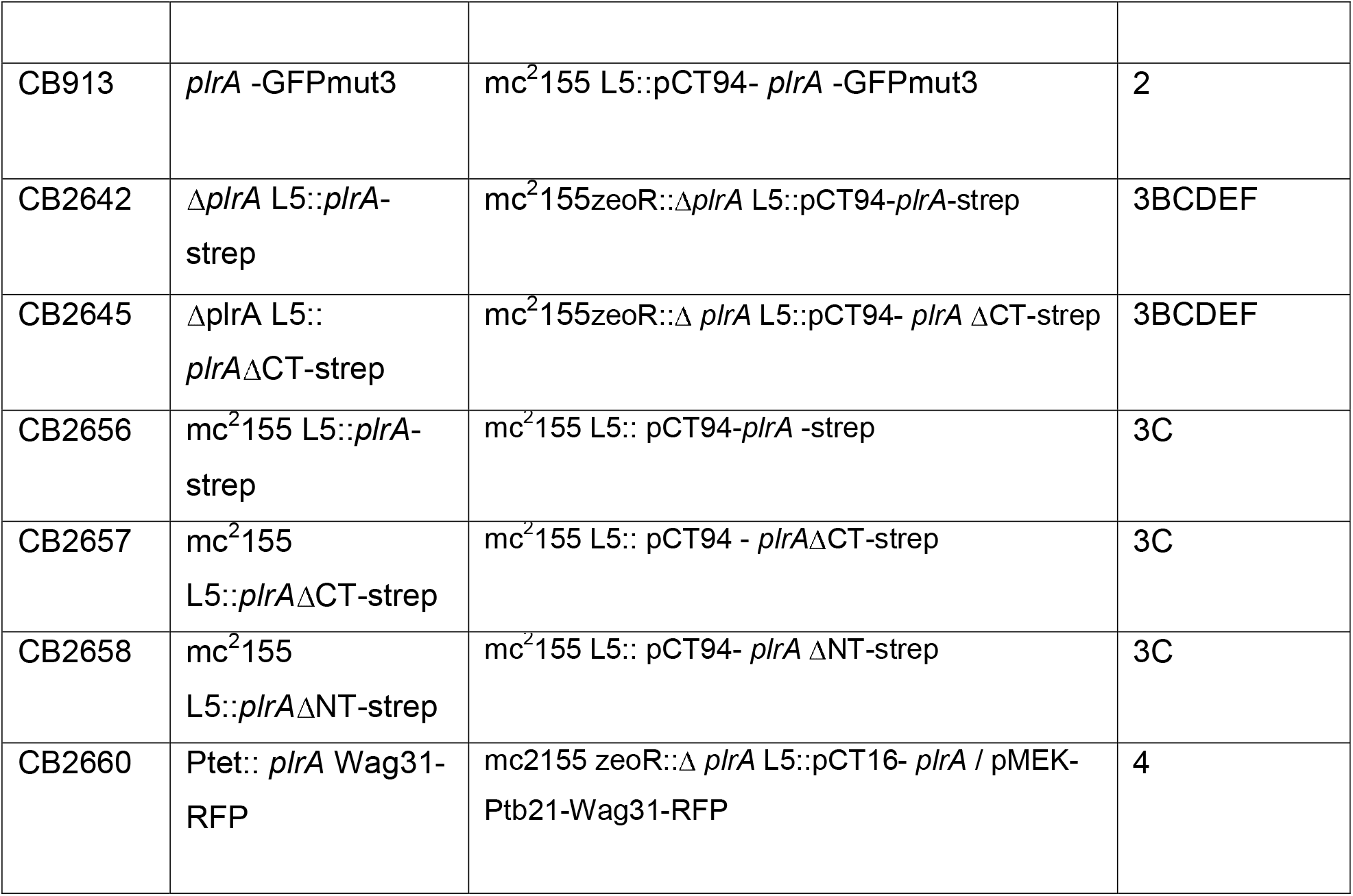
Strains.

| Strains |  |  |  |
| --- | --- | --- | --- |
| Strain # | nickname | genotype | Figure panel |
| CB966 | Ptet:: *plrA* | mc<sup>2</sup>155 Δ *plrA*::zeoR L5::pMC1s- *plrA* | 1 |
| CB913 | *plrA* -GFPmut3 | mc<sup>2</sup>155 L5::pCT94- *plrA* -GFPmut3 | 2 |
| CB2642 | Δ*plrA* L5::*plrA*-strep | mc<sup>2</sup>155zeoR::Δ*plrA* L5::pCT94-*plrA*-strep | 3BCDEF |
| CB2645 | Δ*plrA* L5::*plrA*ΔCT-strep | mc<sup>2</sup>155zeoR::$\Delta$ *plrA* L5::pCT94- *plrA* ΔCT-strep | 3BCDEF |
| CB2656 | mc<sup>2</sup>155 L5::*plrA*-strep | mc<sup>2</sup>155 L5:: pCT94-*plrA* -strep | 3C |
| CB2657 | mc<sup>2</sup>155 L5::*plrA*ΔCT-strep | mc<sup>2</sup>155 L5:: pCT94 - *plrA*ΔCT-strep | 3C |
| CB2658 | mc<sup>2</sup>155 L5::*plrA*ΔNT-strep | mc<sup>2</sup>155 L5:: pCT94- *plrA* ΔNT-strep | 3C |
| CB2660 | Ptet:: *plrA* Wag31-RFP | mc2155 zeoR::$\Delta$ *plrA* L5::pCT16- *plrA* / pMEK-Ptb21-Wag31-RFP | 4 |

**Supplemental Table 2.** Plasmids. *If a published vector was used unaltered, it is indicated with an * in the “Ref for parent/vector” column.

| Plasmids. |  |  |  |
| --- | --- | --- | --- |
| strain # | Plasmid name | Used in strains | Ref for parent/ vector* |
| CB964 | pMC1s- *plrA* | CB966,<br>CB2321 | [53] |
| CB909 | pCT94- *plrA*-GFPmut3 | CB1126 | [54] |
| CB1401 | pCT94-MSMEG_5223-strep | CB2656,<br>CB2642 | [54] |
| CB2636 | pCT94 - 5223- $\Delta$ CT-strep | CB2657,<br>CB2645 | [54] |
| CB2637 | pCT94 - 5223- $\Delta$ NT-strep | CB2658 | [54] |
| CB 1261 | pMEK-Ptb21-Wag31-RFP | CB2660 | [23] |

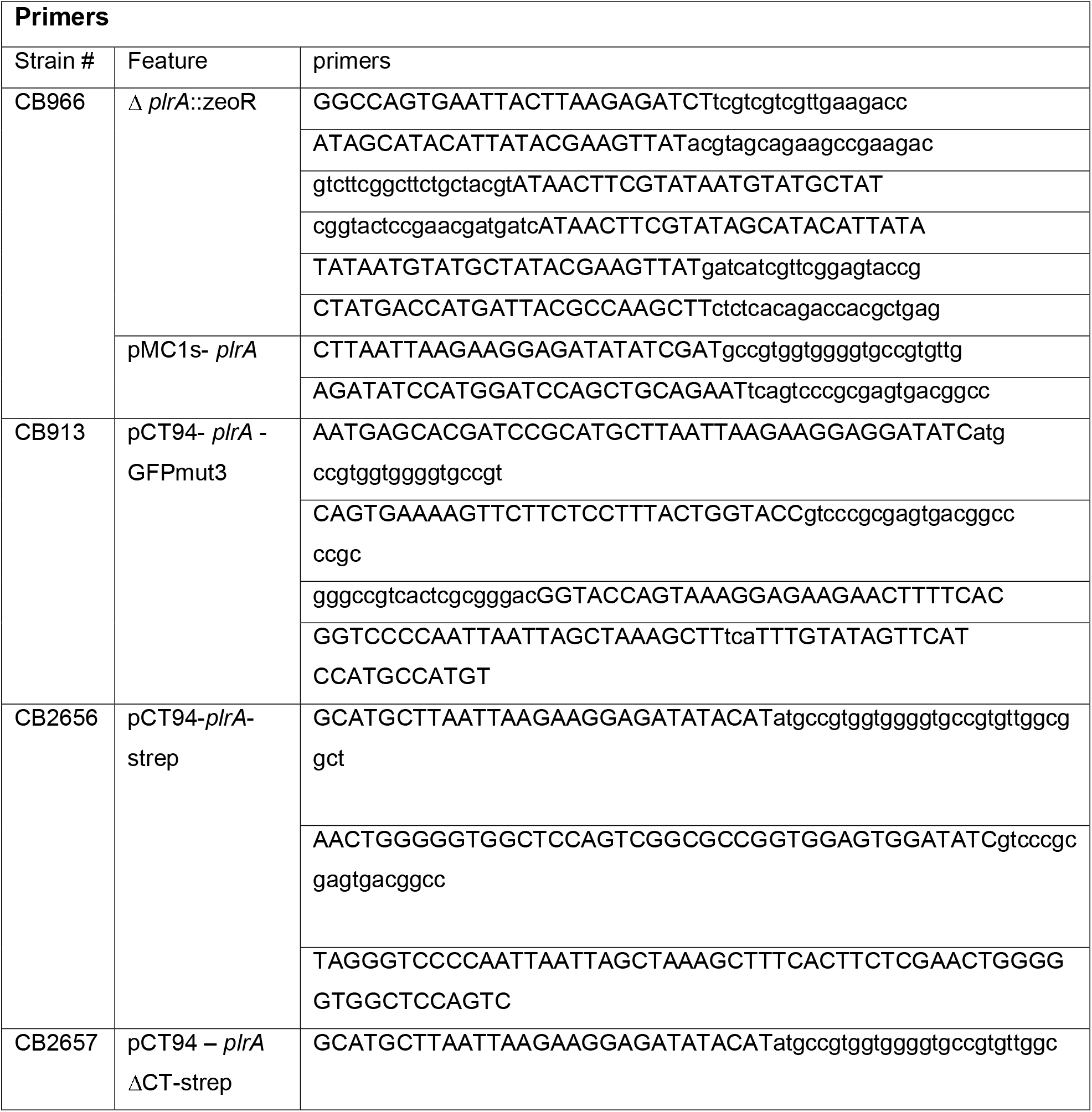
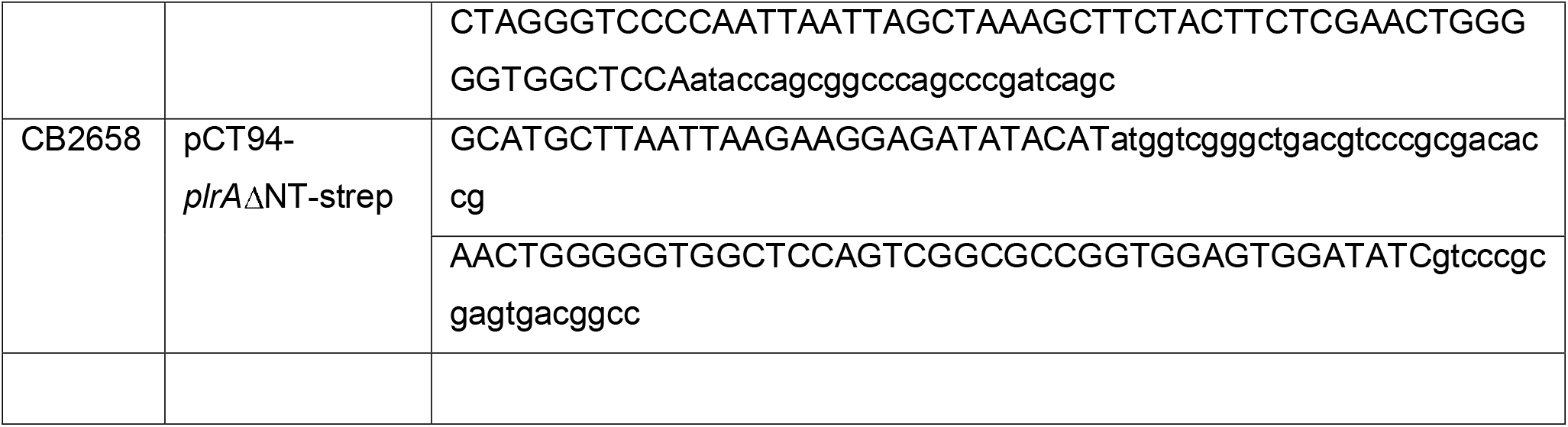

